# MRI of *Capn15* knockout mice and analysis of Capn 15 distribution reveal possible roles in brain development and plasticity

**DOI:** 10.1101/763888

**Authors:** Congyao Zha, Carole A Farah, Vladimir Fonov, David A Rudko, Wayne S Sossin

**Affiliations:** Department of Neurology and Neurosurgery, Montreal Neurological Institute, McGill University, Montreal, Quebec, Canada, H3A 2B4; Department of Biomedical Engineering, McGill University, Montreal, Quebec, Canada

**Author notes:** Equally contributed (Congyao Zha, Carole A Farah). Corresponding author: Wayne S Sossin.

**Keywords:** Calpains, hippocampus, brain development, Purkinje neurons, small animal MRI, brain size, SOL

## Abstract

The Small Optic Lobe (SOL) family of calpains are intracellular cysteine proteases that are expressed in the nervous system and play an important role in neuronal development in both *Drosophila*, where loss of this calpain leads to the eponymous small optic lobes, and in mouse and human, where loss of this calpain leads to eye anomalies. Some human individuals with biallelic variants in CAPN15 also have developmental delay and autism. However, neither the specific effect of the loss of the Capn15 protein on brain development nor the brain regions where this calpain is expressed in the adult is known. Here we show using small animal MRI that mice with the complete loss of Capn15 have smaller brains overall with larger decreases in the thalamus and subregions of the hippocampus. These losses are not seen in *Capn15* conditional KO mice where Capn15 is knocked out only in excitatory neurons in the adult. Based on β-galactosidase expression in an insert strain where lacZ is expressed under the control of the *Capn15* promoter, we show that Capn15 is expressed in adult mice, particularly in neurons involved in plasticity such as the hippocampus, lateral amygdala and Purkinje neurons, and partially in other non-characterized cell types. The regions of the brain in the adult where Capn15 is expressed do not correspond well to the regions of the brain most affected by the complete knockout suggesting distinct roles of Capn15 in brain development and adult brain function.

## Introduction

Calpains are intracellular cysteine proteases that were first discovered in rat brain (Guroff, 1964). There are four conserved families of calpains: Classical, PalB (phosphatase mutants: loss in activity at alkaline pH, but normal or increased activity at acidic pH), Transformer (Tra), and Small Optic Lobe (SOL) with the classical being the best characterized family. All calpain isoforms have a conserved catalytic domain and each calpain family has unique domains. Of interest to the present study, SOL calpains (Capn15 in vertebrates) have an N-terminal zinc finger domain that binds polyubiquitin (Hastings et al., 2018) and a C-terminal SOL homology domain (Hastings et al., 2017;Zhao et al., 2012). The most ancient family of calpains is PalB which is expressed in Fungi, followed by SOL which is expressed in the earliest metazoans, while Tra and classical calpains diverged from PalB in early pre-bilaterians (Hastings, et al., 2017). Thus, of the four families of calpains present throughout animals, SOL is the most diverged (Hastings, et al., 2017).

Calpains play diverse roles in cellular physiology (Bertipaglia and Carafoli, 2007;Ono and Sorimachi, 2012). In the nervous system, degradation of repressors of plasticity by classical calpain, Capn1, in the hippocampus is important for the induction of synaptic plasticity associated with memory (Briz and Baudry, 2017;Khoutorsky et al., 2013;Shimizu et al., 2007). Another distinct classical calpain, Capn2, plays an opposite role in limiting plasticity (Liu et al., 2016;Wang et al., 2014). Furthermore, *Capn2* deletion from birth results in embryonic lethality (Dutt et al., 2006) demonstrating multiple distinct roles for this specific calpain isoform in development and in the adult. However, while the role of the classical calpains has been well characterized, the role of the other calpain family members in vertebrates is less clear.

SOL calpain was first identified in the fruit fly *Drosophila* (Delaney et al., 1991;Fischbach and Heisenberg, 1981) where loss of the SOL calpain lead to a 50% reduction in the volume of *Drosophila* optic lobes (Fischbach and Heisenberg, 1981). In *Aplysia*, expression of a dominant negative catalytically inactive SOL calpain specifically prevented a form of non-associative long-term facilitation (Hu et al., 2017a). However, how SOL calpain is activated and the identity of its substrates remains unknown (Hastings, et al., 2018).

We have recently generated *Capn15* knockout (KO) mice and showed that loss of *Capn15* leads to a lower Mendelian ratio, a smaller weight of weaned *Capn15* KO mice and developmental eye anomalies (Zha et al., 2020). Human individuals with biallelic variants in CAPN15 also have developmental eye anomalies (Zha, et al., 2020), and a subset of these individuals have developmental delays and autism. However, how loss of Capn15 affects brain development and where Capn15 is expressed in the adult are not known.

In the present study, using 7 T pre-clinical MR imaging, we reveal a decrease in whole brain volume of adult *Capn15* KO mice, as well as in specific regions, such as the thalamus and certain hippocampal subregions. We generate a *Capn15* conditional KO mouse (cKO) in which Capn15 is only removed from a subset of excitatory neurons in the forebrain after early brain development. Unlike *Capn15* KO mice, *Capn15* cKO mice have normal brain volumes as determined using 7 T MR imaging, as well as normal weight and lack eye anomalies. We also characterize the distribution of Capn15 in the brain of adult *Capn15* KO mice and show that Capn15 is expressed in adult hippocampal CA1 and CA3 neurons and in Purkinje cells in the cerebellum, neurons particularly implicated in synaptic plasticity.

## Material and methods

### Generation of the *Capn15* conditional KO mouse

The generation of the *Capn15^(lacZ-Neo)^* mouse, the FLOXed *Capn15* mouse and the complete *Capn15* KO mouse has been described (Zha, et al., 2020). To generate a conditional KO line (cKO), the mice were bred with the CaMKIIα-Cre-CRE T29-1 (Tsien et al., 1996) that should only delete exons of *Capn15* after development in forebrain excitatory neurons. The initial breeding was flox/flox x CaMKIIα-Cre-CRE T29-1/WT flox/WT. Only a small number of cKO mice were generated from this cross due to the apparent presence of CRE and *Capn15* on the same chromosome (*Capn15* is on chromosome 17 in mouse, the location of the CRE T29-1 has not been described). These cKO mice were used in subsequential breeding as they had presumably recombined the CRE and the floxed *Capn15* allele on the same chromosome. Specifically, we generated cKO mice by breeding cKO mice flox/flox CaMKIIα-Cre-CRE T29-1/WT with flox/flox mice. The CRE-littermates were used as controls.

### Dissections

Adult mice (3-6 months) were transcardially perfused with ice-cold phosphate-buffered saline (PBS) followed by 4% (wt/vol) ice-cold paraformaldehyde (PFA) in PBS. Brains were post-fixed in 4%PFA for 45min at 4°C, rinsed in PBS, and cryoprotected in 30% sucrose/PBS overnight at 4°C. The following day, brains were embedded in Tissue-Plus™ O.C.T. compound (Fisher Healthcare) and flash frozen in 2-methylbutane chilled in dry ice. The brains were kept at −80°C until further use. For amygdala dissection, mouse brains were removed and frozen on dry ice and kept at −80°C until further use.

The amygdala was dissected from each frozen brain in the cryostat using a neuro punch (0.5 mm; Fine Science Tools). For MRI analysis, adult mice were transcardially perfused with ice-cold PBS followed by 4% (wt/vol) ice-cold PFA in PBS. Brains were then post-fixed in 4% PFA overnight at 4°C, rinsed in 1XPBS, and stored in 1XPBS + 0.05% sodium azide. For brain weight measurement, olfactory bulbs and brain stem were removed before weighing the brains.

### X-gal staining

Sections of 20μm were incubated overnight at 37°C in solution containing 80 mM Na_2_HPO_4_, 20 mM NaH_2_PO_4_, 2 mM MgSO_4_, 5 mM K_3_[Fe(CN)_6_], 5 mM K_4_(Fe(CN)_6_], 0.2% NP-40, 0.1% sodium deoxycholate, and 1.5 mg/ml X-gal. Sections were rinsed in PBS, washed in ethanol (50% for 1min, 70% for 1min, 95% for 1min and 100% for 2X1min), cleared in xylene, and mounted with Permount (Fisher Scientific). 9 adult *Capn15^(lacZ-Neo)^* mice (4 M and 5F) and 2 WT mice were used for X-Gal staining and results were similar for all mice examined.

### Immunohistochemistry

Adult mouse brains were fixed with 2% PLP solution (PBS containing 2% PFA, 10mM sodium periodate, and 70mM L-lysine) for 3 h at 4°C and P3 mouse brains were fixed with 2% PLP solution for 1 hr at 4°C. Brains were cryoprotected in 30% (wt/vol) sucrose in PBS and embedded in OCT (Fisher Healthcare). Brains were frozen on dry ice and cryosectioned (14-μm-thick coronal sections). Sections were first incubated in blocking solution containing 5% (vol/vol) goat serum (or donkey serum for goat antibodies), 0.1% (vol/vol) Triton-X, and 0.5mg/ml BSA in PBS for 1.5 h. Brain sections and eye sections were incubated with primary antibodies in the blocking solution overnight at 4°C. After rinsing five times with the blocking solution for 5 min, sections were incubated with secondary antibodies in the blocking solution for 1 h at room temperature. DNA was labeled with Hoechst (1:2000) for 3 min and the sections were mounted using fluorescence mounting media (Dako, S3023). All images were taken using a Zeiss Observer Z1 fluorescent microscope using a 10X objective. For the primary antibody, we used chicken polyclonal antibody directed against β-galactosidase (β-gal; abcam, dilution 1:1000), rabbit polyclonal antibody directed against Purkinje cell protein 4 (PCP4; Sigma, dilution 1:400), rabbit polyclonal antibody raised to Capn15 (Zha, et al., 2020), goat polyclonal antibody directed against Brn3a (Santa Cruz, dilution 1:500), goat polyclonal antibody directed against ChAT (Millipore Sigma, dilution 1:100), rabbit polyclonal antibody directed against GFAP (Abcam, dilution 1:500), rabbit polyclonal antibody directed against Iba-1 (FUJIFILM Wako, dilution 1:500) and mouse monoclonal antibody directed against GAD67 (Millipore Sigma, dilution 1:500). For the secondary antibody, we used goat antichicken secondary antibody conjugated to Alexa Fluor 488, goat anti-chicken secondary antibody conjugated to Alexa Fluor 594, donkey anti-goat IgG secondary antibody conjugated to Alexa Fluor 488, goat anti-rabbit IgG secondary antibody conjugated to Alexa Fluor 488, goat anti-rabbit IgG secondary antibody conjugated to Alexa Fluor 568 and goat anti-mouse IgG secondary antibody conjugated to Alexa Fluor 568 (Invitrogen, dilution 1:500). Overall, experiments were performed on 9 *Capn15^(lacZ-Neo)^* brains and 2 WT brains. All staining were repeated from at least three separate *Capn15^(lacZ-Neo^* mice with the exception of Iba-I staining from two separate animals and GFAP staining which used only one animal.

### Quantification of cell number

Neurons in the CA1 area of the hippocampus were counted manually in 2-7 Nissl-stained sections from 3 WT and 3 *Capn15* KO mice that were previously used for MRI imaging. The hand free tool was used to draw an area in the CA1 region of the hippocampus, and the number of neurons was quantitated per mm^2^ for each section and then averaged for each mouse by an observer blind to the genotype of the animal. The number of cells in PCP4-labeled cerebellar sections were counted manually in 8-10 Nissl sections through the cerebellum. Density was calculated as the number of cells per 100 μM as Purkinje cells are mainly contained in a single row of cells. The density of cells was calculated for WT and *Capn15* KO animals by an observer blind to the genetic status of the animal.

### Western blotting

Brains were homogenized manually in lysis buffer containing 25 mM Tris-HCl (pH 7.4), 150 mM NaCl, 6 mM MgCl_2_, 2 mM EDTA, 1.25% NP-40, 0.125% SDS, 25 mM NaF, 2 mM Na_4_P_2_O_7_, 1 mM dithiothreitol (DTT), 1 mM phenylmethylsulfonyl fluoride (PMSF), 20 mg/ml leupeptin, and 4 mg/ml aprotinin. Before loading, 5X sample buffer was added to the lysate and samples were incubated at 95°C for 5min. Proteins were resolved by SDS-PAGE on Bis-Tris gel and transferred to nitrocellulose membrane (Bio-Rad). The blots were blocked in TBST (TBS + 0.1% Tween) containing 4% skim milk for 30min at room temperature and then incubated with primary antibodies overnight at 4°C. After washing 3 times with TBST, the blots were incubated with HRP-conjugated secondary antibodies for 1 hour at RT and washed again 3 times in TBST. The Western Lightning Plus-ECL kit (NEL103001EA; PerkinElmer LLC Waltham, MA USA) was used as per manufacturer’s instructions to detect protein bands. The primary antibody used was a homemade rabbit anti-Capn15 antibody (1:1000) raised against the C-terminus of Capn15 (Zha, et al., 2020),. The secondary antibody was horseradish peroxidase-conjugated goat anti-rabbit secondary antibody (1:5000). Antibodies were diluted in Tris buffered saline with Tween containing 4% skim milk powder.

### Eye phenotype quantification

Mice eyes were examined and grouped as follows: seems normal, obvious cataract, small eye and no eye. This categorization was performed for both eyes of each mouse. The analysis was performed without the knowledge of the genotype of the mice.

### Quantification of immunoblotting

Immunoblots were scanned and imaged using the public domain Image J program developed at the U.S. National Institute of Health (https://imagej.nih.gov/ij/). We calibrated our data with the uncalibrated optical density feature of NIH image, which transforms the data using the formula 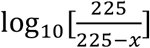, where x is the pixel value (0–254). We find that with this correction and including the entire band (which expands near saturation), values are linear with respect to amount of protein over a wide range of values (Nakhost et al., 1998). We used the Ponceau image for each gel to normalize the amount of SOL calpain to total protein loaded.

### Image Acquisition

Magnetic resonance imaging was performed using the 7 T Bruker Pharmascan (Bruker Biosciences, Billerica, MA) ultra-high field MRI system of the McConnell Brain Imaging Centre. For imaging, brains were housed in a cylindrical container and immersed in an MR-invisible fluorinated solution, FC-40 (Sigma Aldrich, St. Louis, Missouri), to remove the background MRI signal. For MRI radiofrequency excitation and reception, a 2.3 cm inner diameter volume resonator was utilized. The imaging protocol included a sagittal orientation, 3D steady-state free precession MRI sequence with an echo time (TE) of 5 milliseconds, repetition time (TR) of 10 milliseconds, receiver bandwidth of 50 kHz and excitation pulse flip angle of 30 degrees. The image acquisition matrix was selected to achieve an isotropic voxel resolution of 100 μm^3^. The specific sagittal orientation imaging field of view was 2.3 × 1.7 × 1.7 cm^3^. 31 signal averages were collected to improve image signal-to-noise ratio, leading to a total scan time of approximately six hours and 60 seconds for the full MRI histology scan. For image reconstruction, a trapezoidal filter was applied to the complex image data along the first phase encode direction to reduce subtle effects of Gibbs ringing.

### Image Processing and Statistical Analysis

We employed an image registration-based method to investigate anatomical brain volume differences between WT, cKO and KO mice. Specifically, an automated, image intensity-based, affine registration using Elastix tool (Klein et al., 2010) was applied to align all mouse brains to a common coordinate system. During this registration step, it was determined that the KO mouse brains were, on average, 14% smaller than the WT mouse brains. For this reason, scans of all animals were linearly scaled to match the average size of the population. To avoid registration bias caused by differences between our studied population and the mouse template (Kovacevic et al., 2005), a population-specific average was constructed using an algorithm described in Fonov et al (Fonov et al., 2011). This step yielded an average image of all the samples included in our cohort. A mouse brain atlas (Kovacevic, et al., 2005) containing reference regions of interest (ROIs) for the thalamus, amygdala, parieto-temporal cortex and whole hippocampus was next used to provide suitable ROIs for volumetric measurement. More specifically, a non-linear registration (ANTs (Avants et al., 2008)) between the atlas space (Kovacevic, et al., 2005) and our population average was conducted to create an anatomical atlas specific to our mouse population. The deformation fields from the non-linear registration were inverted. The Jacobian determinants of the inverted deformation fields were then calculated to yield estimates of local expansion or contraction at both the ROI and voxel level relative to the population average. We smoothed the Jacobian determinant fields using a Gaussian smoothing kernel with a full width half maximum (FHWM) of 0.2 mm. The choice of the FWHM was based on both image signal-to-noise ratio and the size of the smallest features of interest in the image. We decided to use a FWHM of 0.2 mm as a tradeoff between localization accuracy and noise reduction.

Since we also had a specific interest in volumetric changes in KO mice in the hippocampal sub-fields, a mouse hippocampal sub-field atlas (Badhwar et al., 2013) was non-linearly registered (Avants, et al., 2008) and warped to the population average to measure volumes of the Cornu Ammonis 1 (CA1), CA2, CA3, stratum granulosum and dentate gyrus in the mouse brain images.

We next used a linear model in the R software package (Chambers, 1992; RTeam, 2020) to measure the main effect of volume differences (i) in individual ROIs in KO and cKO mice compared to WT mice and (ii) at the voxel-level in KO and cKO mice compared to WT mice. All statistical analyses were corrected for multiple comparisons, utilizing the false discovery rate technique (Genovese et al., 2002) with a 5% threshold. Voxel-based analysis and generation of figures was performed using RMINC package (Lerch J. et al., 2017). Measurements from the left and right hemisphere were always averaged for volume measurements.

## Results

### Generation of a conditional *Capn15* KO mouse

To assess the role of *Capn15* in the adult, we generated *Capn15* conditional KO mice (cKOs) as described in the methods and a CaMKIIα-Cre line was used to this end. CaMKIIα-Cre transgenic mice have the mouse calcium/calmodulin-dependent protein kinase II alpha promoter driving Cre recombinase expression in a subset of excitatory neurons in the forebrain including the hippocampus and the amygdala and CRE is not expressed until after P14 (Sonner et al., 2005). As shown in Figs.1A-D, Capn15 protein levels are significantly decreased in homogenates from *Capn15* cKO adult mice amygdala (\*\**p*=0.002, Student’s t-test; n=4 cKO, 3 WT) and hippocampus (*p=0.01, Student’s t-test; n=4 cKO, 4 WT) indicating successful disruption of the *Capn15* gene in cKO mice. Complete *Capn15* KO mice have reduced weight and eye anomalies (Zha, et al., 2020). In contrast, *Capn15* cKO mice showed no weight difference compared to WT mice (*p*=0.16, Student’s t-test; n=26 for the WT group composed of *Capn15* cKO Cre-littermates and n=15 for the *Capn15* cKO group; Fig.1E) and no eye deficits (Fig.1F, n=20 *Capn15* cKO, n=20 WT).

**Figure 1.**
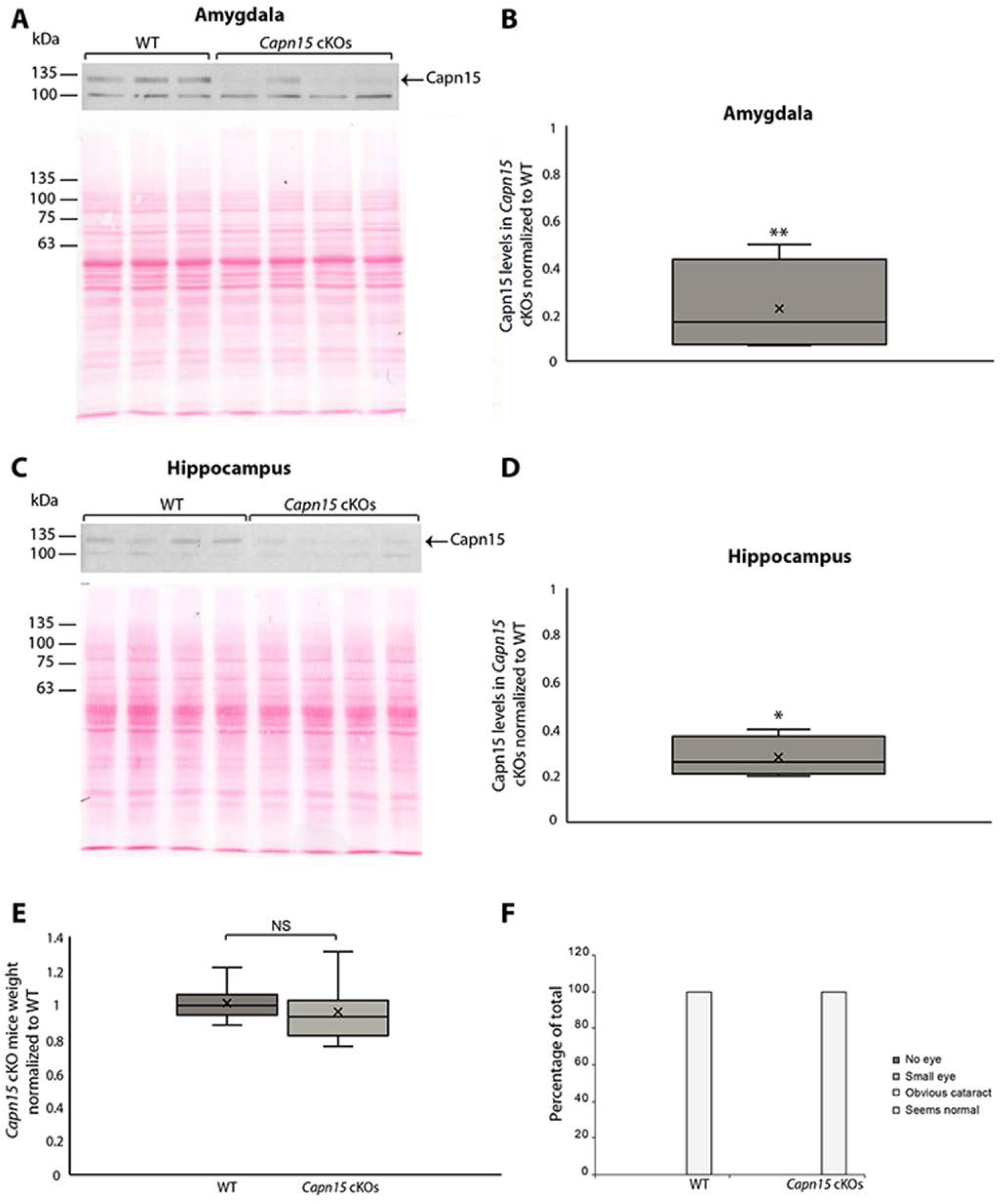
*Capn15* cKO mice have reduced Capn 15 levels, but normal weight and eyes. Amygdala **(A)** and hippocampus **(C)** were dissected and homogenized and SDS-PAGE and western blotting were performed as described in Methods. **(B)** Box and whisker plot showing that Capn15 protein levels are significantly decreased in *Capn15* cKO adult mice amygdala, **(D)** Box and whisker plot showing that Capn15 protein levels are significantly decreased in *Capn15* cKO adult mice hippocampus. **(E)** Animals were weighed after weaning. Average weight of *Capn15* cKO was normalized to average weight of the WT mice. **(F)** The percentage of mice having eye deficits between WT and *Capn15* cKO mice. Mice were grouped in 4 categories: Seems normal, obvious cataract, small eye and no eye and this scoring was performed on both left and right eye for each mouse. All of WT and *Capn15* cKO mice eyes scored normal.

### *Capn15* KO mice have smaller brains

In *Drosophila*, loss of the SOL calpain was mainly reported to cause deficits in the optic lobe (Fischbach and Heisenberg, 1981). However, it is not clear how extensively other brain regions were analyzed. To use a non-biased screen to determine which brain regions are affected by the loss of *Capn15*, we applied whole brain 7 T MRI on 14 WT (9M, 5F), 8 *Capn15* KO (4M, 4F) animals and 5 *Capn15* cKO (5M). There was no difference in whole brain volume between male and female mice for either the WT (*p*=0.97) or the *Capn15* KO groups (*p*=0.83). Accordingly, males and females were grouped together for statistical analysis. There was a highly significant, 14% decrease in whole brain volume in the *Capn15* KO animals (*p*=7.32e-7) compared to the WT and cKO animals (Fig.2A). Measurement of the brain weight of WT and *Capn15* KO animals confirmed that the brains of *Capn15* KO mice were significantly lighter than WT brains, similar to the amount measured by 7 T MRI (0.389g ± 0.002 for the WT group composed of *Capn15* KO littermates, n=14; 0.334g ± 0.002 for the *Capn15* KO group, n=12;*p*=7.28e-14, Student’s t-test, SEM).

**Figure 2.**
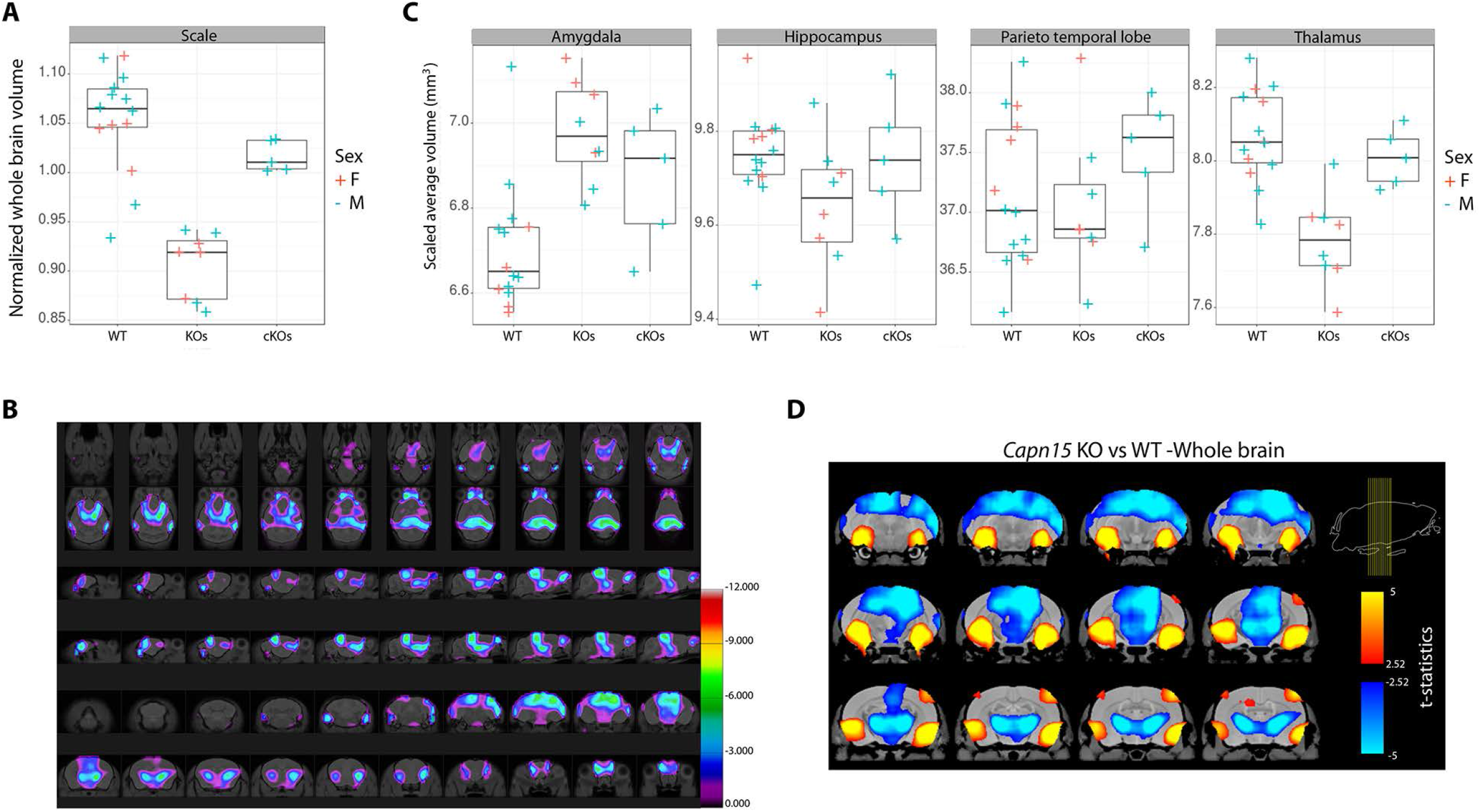
Whole-brain visualization of significant decreases in brain structure volume in *Capn15* KO mice relative to cKO and WT mice. **(A)** Box and whisker plots showing scaling factor (see Methods, Fonov et al, 2011) used for each animal to compensate for differences in overall brain size. This scaling factor was applied to all structures shown in Figure 2 and 3. **(B)**.The top two rows correspond to 20 axial sections through the mouse brain, the middle two rows correspond to 20 sagittal sections through the brain and the bottom two rows correspond to 20 coronal sections. Each section has a thickness of 100 micrometers. Purple/blue/green colour maps overlaid on the mouse brain images identify logio(p) values at the voxel-level. The colour maps specifically show regions where a significant brain volume decrease was observed for KO mice relative to WT mice. All the logio(p) values were calculated with a general linear model. Multiple comparisons correction was carried out using the false discovery rate method with a q-threshold of 0.05. **(C)** Structure-specific brain volume differences among WT, KO and cKO mice Box and whisker plots showing volume differences of different brain regions among WT, KO and cKO mice. All anatomical regions in each brain were scaled based on scaling factor shown in A) to compensate for the overall brain size differences (See methods) **(D)** Magnified coronal views through the mouse mid-brain displaying regional volume decrease (blue colours) and increase (orange) for KO mice compared to WT mice. Each slice through the mid-brain has a thickness of 100 micrometers. The specific position of each slice along the anterior-posterior axis of the mouse brain is shown, overlaid on a sagittal brain slice, in the top right-hand corner of the figure. The colours represent t-statistical values calculated with a general linear model. Multiple comparisons correction was carried out using the false discovery rate method with a q-threshold of 0.05.

### *Capn15* KO brains have specific decreases in thalamic and hippocampal region volumes

We next examined decrease in volume in specific brain regions in *Capn15* KO, *Capn15* cKO and WT mice after first compensating for the overall decrease in brain size of *Capn15* KO animals (See Methods). The most significant decrease in volume were observed in the hemisphere-averaged thalamic volumes of *Capn15* KO animals compared to the corresponding WT animals (*p*=1.28e-5; Figs.2B and 2C). There was no difference in thalamic volume between *Capn15* cKO and WT animals (*p*=0.35; Figs.2B and 2C). Atlas-based region of interest analysis revealed an increase in the amygdala volume of *Capn15* KO mice compared to WT mice (*p*=0.003, Figs.2B and 2C). However, there was no corresponding difference between *Capn15* cKO and WT mice (*p*=0.42). As well, there were no differences in volume of the parieto-temporal cortex between either *Capn15* KO or cKO mice and corresponding WT mice (Figs.2B and 2C). We also used voxel-based morphometry to measure local differences in volume between *Capn15* KO and WT mice. This was specifically performed to leverage the high spatial resolution (100 micron isotropic voxels) of the anatomical mouse brain images to carry out an unbiased test of local, voxel-specific volume differences between KO, cKO and WT mice. In essence, unlike ROI-based analysis where the mean volume difference between genotypes is calculated inside an anatomically defined regions, voxel-based morphometry evaluates volume differences at a very small scale without predefined brain structural boundaries. Twelve representative slices having locally decreased volume of *Capn15* KO brain compared to WT brain (blue colours) and the corresponding locally increased volume (yellow/orange colours) are presented in Fig.2D. Volume decreases in *Capn15* KO mouse brain are notable (light blue colours) in the hippocampus, superior parietal cortex and thalamus.

Additional attention in our high resolution, MRI-based volumetric analysis was given to the hippocampus due to the possible role of Capn15 in synaptic plasticity (Hu, et al., 2017a;Hu et al., 2017b). An image of the hippocampal sub-field atlas we employed for sub-field volumetry is shown in Fig.3A. We did not observe differences in overall hippocampal volume between either *Capn15* KO or *Capn15* cKO mice and the corresponding WT animals. However, voxel-level differences in hippocampal volume were observed between *Capn15* KO and WT mice when we restricted our linear model statistical analysis to regions inside the hippocampal sub-fields (Figs.3A and 3B). In particular, there were voxel decreases in hippocampal volume for mid-line slices of the *Capn15* KO mice brain compared to the WT mice. Examination of the hippocampal subfields using region of interest-based analysis identified decreases in the CA1 (*p*=0.001), the CA2 (*p*=0.002), the dentate gyrus (*p*=0.05) and the stratum granulosum (p=0.03) volume of *Capn15* KO mice when compared to WT mice (Figs.3A and 3B). Notably, such volumetric differences were not observed in CA3. It is possible that localized significant changes in volume are taking place in sub-regions of the CA3 area but the hippocampal atlas label includes both regions that increase and regions that decrease in volume. Therefore, the box plot does not show statistically significant aggregate volume change in the CA3 label. As well, no differences in hippocampal sub-field volume were observed between *Capn15* cKO and WT mice. We further performed a voxel-level statistical analysis of contraction/expansion inside the hippocampus to test the hypothesis that within the sub-fields of the mouse hippocampus volume differences exist between KO, cKO and cKO mice. As shown in Fig.3C, volume decreases in *Capn15* KO mouse brain are notable (light blue colours) in the CA1, CA2, dentate gyrus and stratum granulosum whereas increases (orange) are noted in the CA3 area.

**Figure 3.**
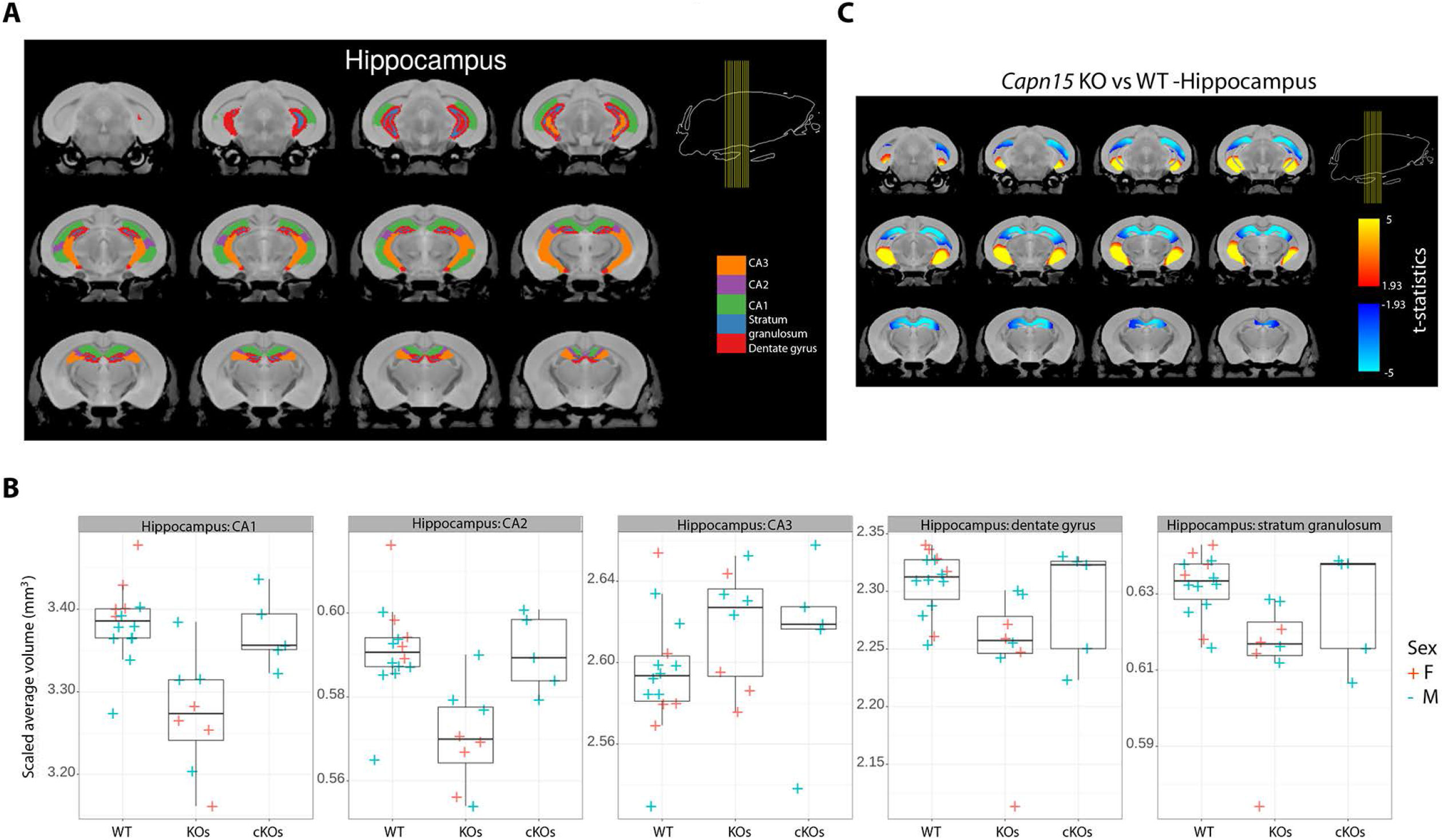
Hippocampal subfield-specific volume changes in *Capn15* KO compared to cKO and WT mice. **(A)** A hippocampal sub-field atlas (Badhwar, et al., 2013) was applied to evaluate volumetric changes in the hippocampal subfields. The sub-field atlas includes region of interest labels demarcating the dentate gyrus (red), stratum granulosum (blue), CA1 (green), CA2 (purple) and CA3 (orange). **(B)** Box and whisker plots showing hippocampal subfield-specific volume differences between WT, KO and cKO mice. All anatomical regions in each brain were scaled based on scaling factor shown in A) to compensate for the overall brain size differences (See methods) **(C)** Voxel-level t-statistical parametric maps showing areas of volumetric contraction (blue colours) or expansion (orange colours) for *Capn15* KO mice compared to WT mice inside the body of the hippocampus. The t-statistical values were calculated with a general linear model. The false discovery rate method was applied for multiple comparisons correction with a q-threshold of 0.05. Each slice through the mid-brain has a thickness of 100 micrometers. The specific position of each slice along the anterior-posterior axis of the mouse brain is shown, overlaid on a sagittal brain slice, in the top right-hand corner of the figure.

### Capn15 is expressed in areas important for brain plasticity in the adult mouse

We have previously described *Capn15* KO mice (Zha, et al., 2020) and took advantage of the initial mouse strain with a reporter gene *lacZ* that is under the control of the *Capn15* promoter to characterize distribution of Capn15 in developing brain (the *lacZ* reporter is deleted in the generation of the Floxed Capn15 mouse (Zha, et al., 2020)). Capn15 was mainly enriched in the mantle zone in E12 embryos, in the subventricular zone, immediately next to the ventricular zone in E18 embryos and was ubiquitously expressed in the brain in P3 mice (Zha, et al., 2020). We also noted that levels of Capn15, measured using immunoblots of whole brains, notably decreased in the adult, but the areas in the brain that retained Capn15 expression were not determined (Zha, et al., 2020). To expand on these results, we examined Capn15 distribution in adult *Capn15^(lacZ-Neo)^* heterozygous mice using X-gal staining. As shown in Fig.4, Capn15 is still expressed in specific sets of neurons in the mature brain, including the hippocampus, amygdala, Purkinje neurons, and cortex. A closer look at the hippocampal region shows an unequal distribution of staining. Excitatory pyramidal neurons in CA1 and CA3 and some cells located in stratum radiatum are stained. However, a gap near the expected position of CA2 was seen and in the dentate gyrus, Capn15 is expressed at low levels in the subgranular zone and showed the strongest signal in the molecular layer. This region contains mainly interneurons such as molecular layer perforant pathway cells (MOPP) (Li et al., 2013) and neurogliaform cells (Armstrong et al., 2011) as well as astrocytes (Pilegaard and Ladefoged, 1996).

**Figure 4.**
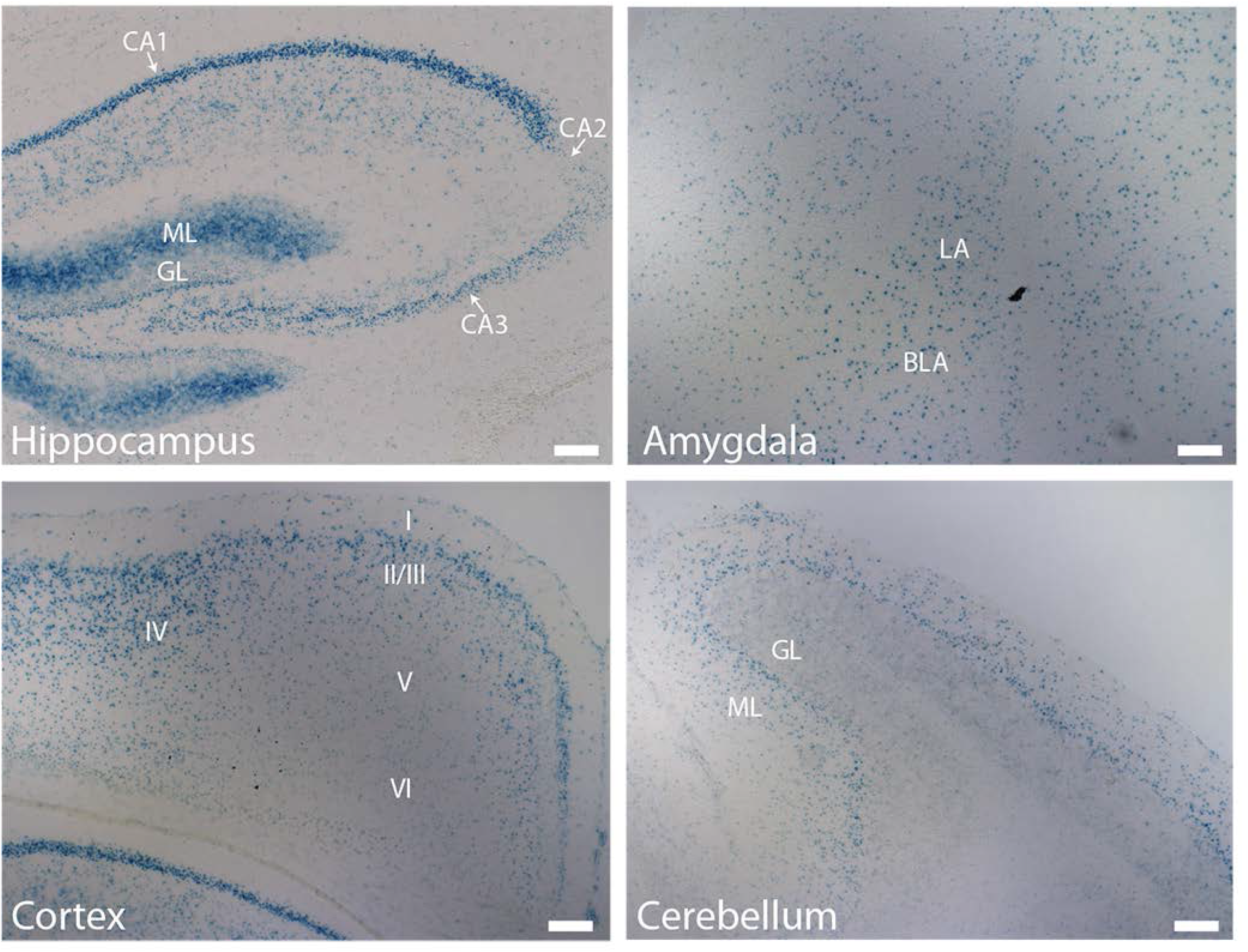
Capn15 distribution in *Capn15^(lacZ-Neo)^* heterozygous mice brain. X-gal staining of coronal sections from adult *Capn15^(lacZ-Neo)^* heterozygous mice hippocampus, amygdala, cortex and cerebellum are shown. Scale bar is 100μm. LA, lateral amygdala; BLA, basal lateral amygdala; GL, granular layer; ML, molecular layer.

To confirm the result obtained with X-gal staining, we stained sections from adult WT and *Capn15* KO mice brains with our home-made antibody to Capn15 (Zha, et al., 2020). Unfortunately, this antibody lacked specificity in immunohistochemistry as staining showed no difference between WT and KO animals and thus, could not be used to this end. Thus, we took advantage of the *lacZ* reporter and stained sections from adult *Capn15-lacZ* mice with an anti β-galactosidase (β-gal) antibody as well as an antibody directed against the CA2 marker PCP4 (Lein et al., 2005) to confirm the lack of Capn15 in CA2 seen with X-gal. DNA was labeled with Hoechst. The β-gal immunoreactivity appeared as punctate spots (Fig.5), perhaps due to aggregation or specific sub-cellular localization of the enzyme in neurons. To confirm β-gal antibody specificity, we costained sections from WT mouse brain with the β-gal antibody and Hoechst. As shown in Fig.5 (top), a weak diffuse signal was observed in WT mice with no punctate spots indicating specificity of the punctate staining in *Capn15-lacZ* mice. Moreover, our result confirmed the absence of Capn15 from the CA2 area (Fig.5 bottom).

**Figure 5.**
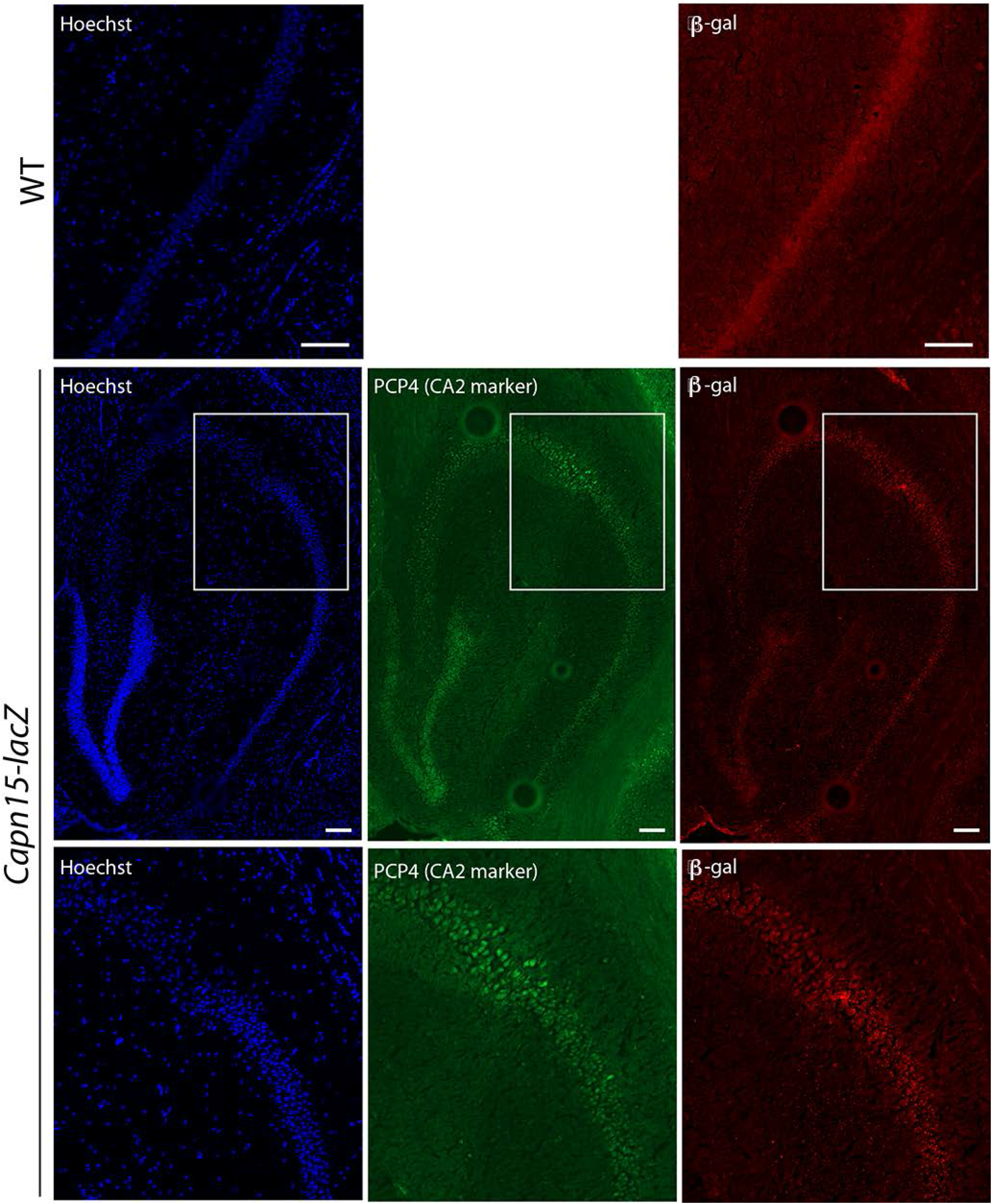
*Capn15* distribution in the hippocampus. The top panel shows the lack of punctate β-gal staining in WT mice. The middle panel shows distribution of a CA2 marker (PCP4) and that of β-gal in a section from the hippocampus from an adult *Capn15-lacZ* brain. The rabbit polyclonal anti-PCP4 and the chicken polyclonal anti-β-gal antibodies were revealed using a secondary anti-rabbit conjugated to Alexa Fluor 488 and a secondary anti-chicken conjugated to Alexa Fluor 594 respectively. DNA was labeled with Hoechst. Capn15 seems to be enriched in all hippocampal areas except in CA2 as illustrated in the magnified inset (boxed region in middle panel) shown in the bottom panel. Scale bars are 100μm.

When examining the staining with the β-gal antibody, we noted a striking difference between the punctate spots in the pyramidal neurons and in the molecular layer. This was most obvious comparing staining in CA3 neurons and the molecular layer in the same sections (Fig.6A). The punctate spots in the molecular layer appeared smaller, more numerous and less associated with nuclei (Fig.6A). Notably, if the X-gal reaction was ended at earlier time points, the molecular layer staining was less intense (Fig.6B), suggesting a lower, but broader level of expression in this area that took longer to saturate. The lower expression of Capn15 in the molecular layer is consistent with the large decrease in overall Capn15 protein observed from immunoblots of adult hippocampal tissue, as the CamKII Cre should not be expressed outside the excitatory neurons in the hippocampus.

**Figure 6.**
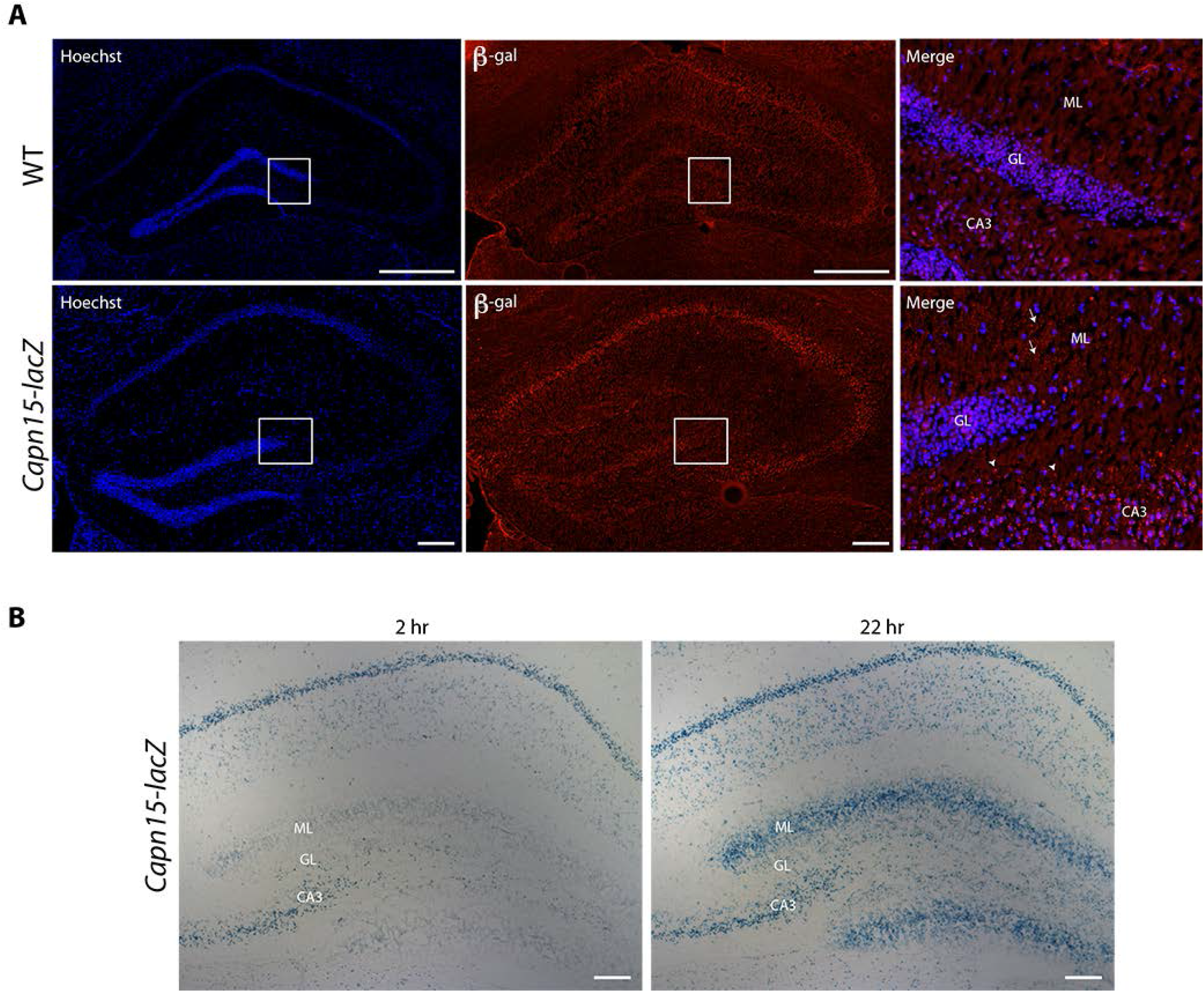
Distinct staining of β-galacatosidase (β-gal) in the CA3 region and the molecular layer. **(A)** The distribution of β-galacatosidase (β-gal) is shown in hippocampal sections from adult *Capn15-lacZ* brains. The chicken polyclonal anti-β-gal antibody was revealed by using a secondary anti-rabbit conjugated to Alexa Fluor 488. DNA was labeled with Hoechst. Capn15 appears to have a distinct expression pattern in the CA3 area and the molecular layer of the hippocampus. Scale bars for top and bottom images are 500μm and 100μm, respectively. Experiments were done on 5 Capn15-lacZ brains and 2 WT brains with similar results. **(B)** X-gal staining of coronal sections from adult *Capn15^(lacZ-Neo)^* heterozygous mice hippocampus is shown. The X-gal reaction was ended at 2 or 22hr. Scale bar is 100μm. GL, granular layer; ML, molecular layer.

We next investigated if β-gal staining in the molecular layer corresponded to astroyctes, microglia, or GABAergic neurons. As shown in Fig.7, β-gal staining did not colocalize with that of the glial marker GFAP (top) or that of the microglia marker Iba-1 (middle) or with the GABAergic marker GAD67 (bottom). At this point, it is not clear what the basis for this staining is, although it was not seen in WT animals (Fig.6A), and thus requires lacZ expression from the *Capn15* locus.

**Figure 7.**
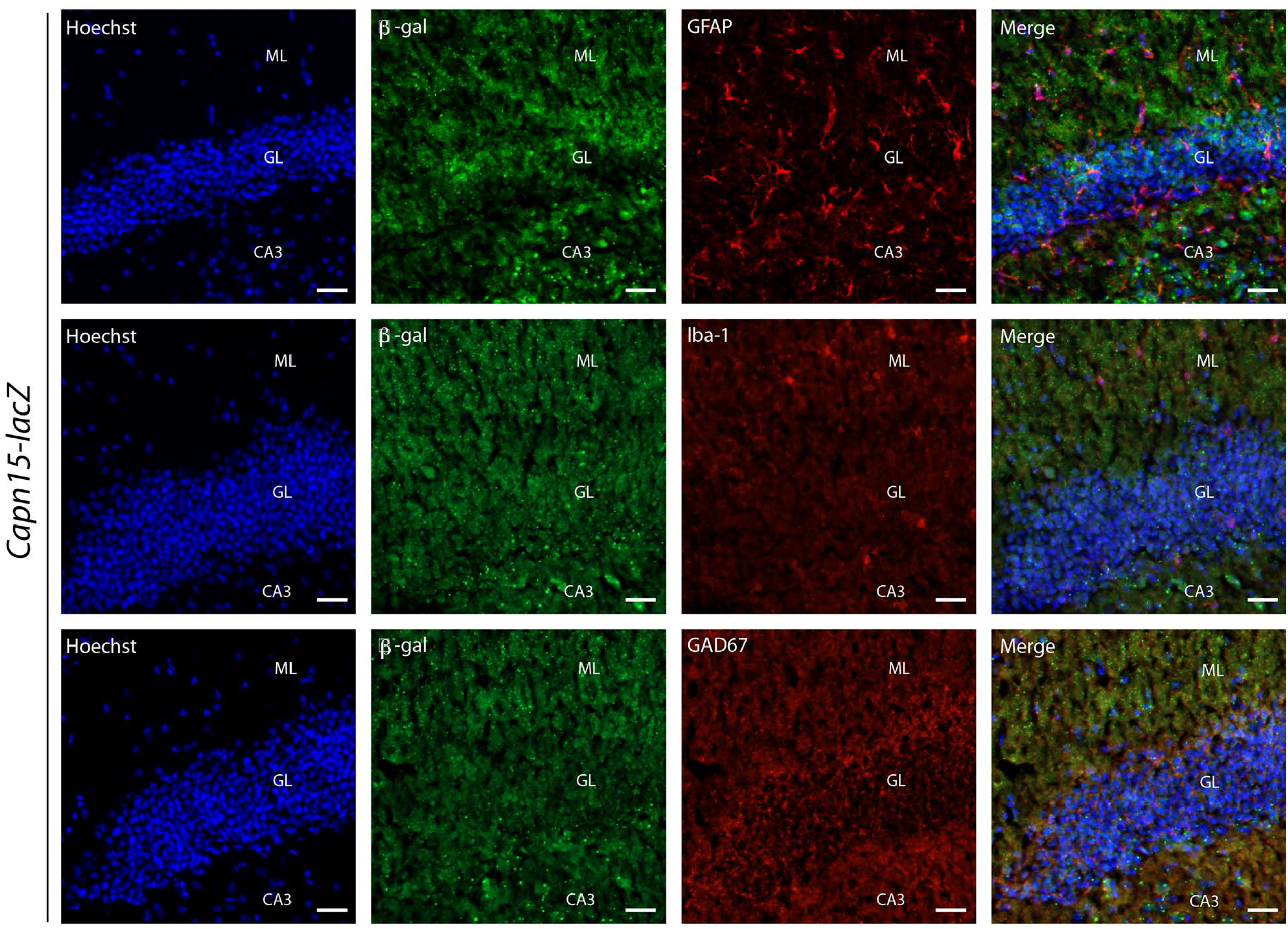
The distribution of β-galacatosidase (β-gal) is shown in hippocampal sections from adult *Capn15-lacZ* brains. From left to right are Hoechst staining, b-gal immunoreactivity, cell-type markers and merged images. From top to bottom the celltype markers are GFAP to label astrocytes, Iba-1 to label microglia and GAD67 to label GABAergic neurons. Scale bar is 25μm. GL, granular layer; ML, molecular layer.

There was a poor correlation between the volume loss in the MRI in the CA2 and CA3 areas of the hippocampus in *Capn15* KO mice and Capn15 expression in these areas in adult *Capn15^(lacZ-Neo)^* heterozygous mice. The CA2 area showed a significant decrease in volume by MRI but did not express Capn15 in the adult, while the CA3 area was relatively larger, but did express Capn15 in the adult. We began to examine whether the loss of volume was correlated with a loss of Nissl-stained cell bodies in the CA1 region. An initial experiment did not detect a difference (2.6 ± 0.3 thousand cells per mm^2^ for the WT group; *n*=3 and 2.7 ± 0.3 thousand cells per mm^2^ for the *Capn15* KO group; *n*=3; SEM, one sample Student’s t-test). However, given the large variance (20%) in the counts in the WT group, we performed a power analysis which suggested that 50 animals would be required to have an 80% chance of detecting a 10% change in cell number and this was not feasible. As these were examined from the same animals on which 7T MRI was performed, we also determined if there was a correlation between CA1 volume (in mm^3^) and CA1 neuronal density derived from cell counting, but no correlation was observed (WT, n=3, KO, n=3, p>0.5)

Staining in the Purkinje neurons was confirmed by staining sections from adult *Capn15-lacZ* mice with the anti-β-gal antibody as well as an antibody directed against PCP4, which was originally described as a Purkinje cell marker (Ziai et al., 1988). DNA was labeled with Hoechst. As shown in Fig.8, Capn15 was enriched in Purkinje cells in the cerebellum. As loss of *Capn15* was linked to neurodegeneration in *Drosophila*, we determined whether the loss of *Capn15* led to a loss of Purkinje cell neurons. We did not detect a difference in the number of Purkinje cells in initial experiments examining *Capn15* KO animals and their WT littermates (3.2 ± 0.4 cells/100μm for the WT group, n=5 compared to 3.2 ± 0.5 cells/100μm for the KO group, *n*=5; SEM, *p*>0.5, Student’s t-test). Again, given the large variation in the number of cells per animal, power analysis suggested over 50 animals would be required to have an 80% chance of detecting a 10% change and this was not feasible.

**Figure 8.**
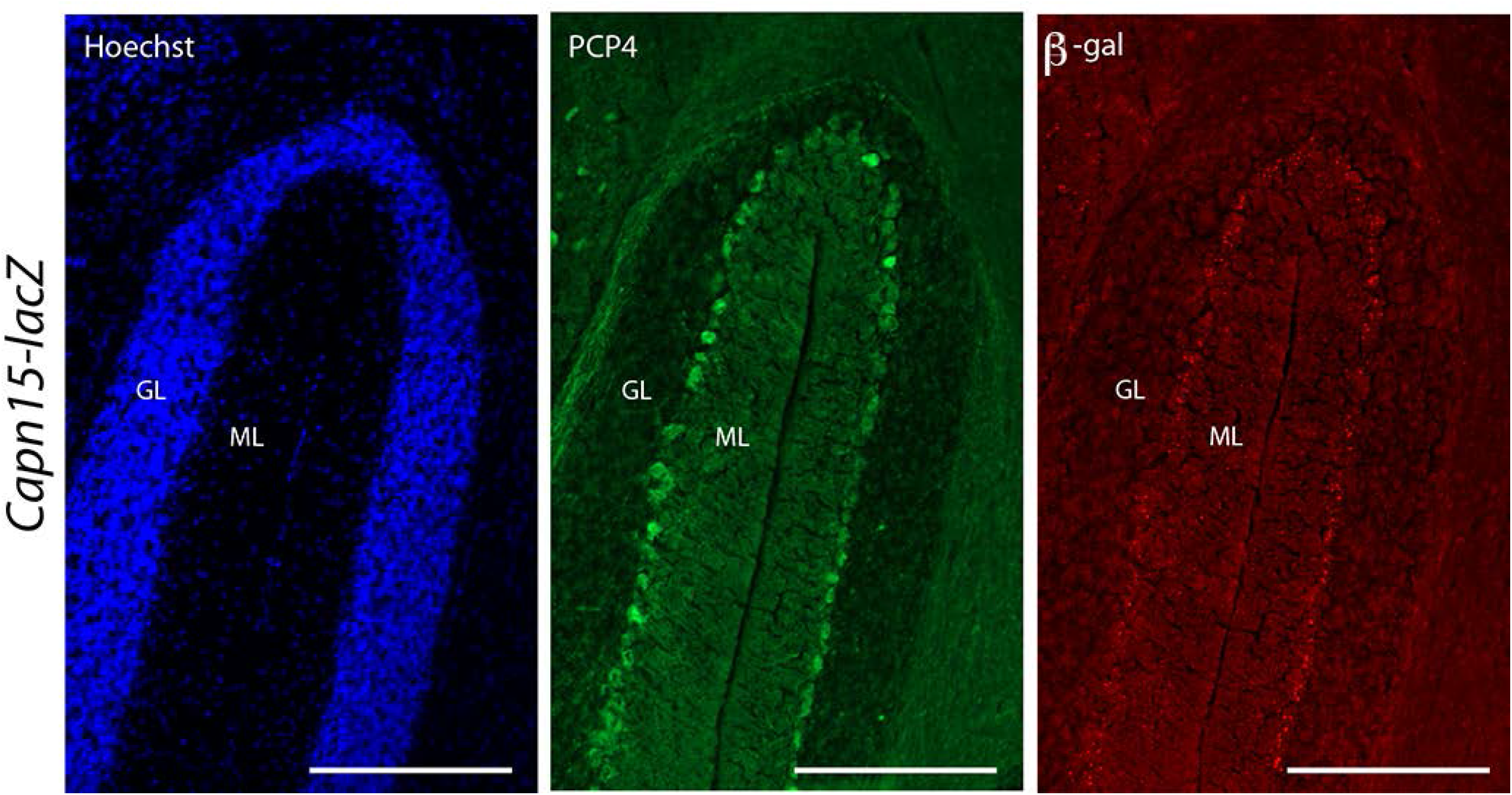
*Capn15* distribution in the cerebellum. The distribution of a Purkinje cell marker (PCP4) and β-galacatosidase (β-gal) is shown in sections from adult *Capn15-lacZ* brains. The rabbit polyclonal anti-PCP4 and the chicken polyclonal anti-β-gal antibodies were revealed by using a secondary anti-rabbit conjugated to Alexa Fluor 488 and a secondary anti-chicken conjugated to Alexa Fluor 594 respectively. DNA was labeled with Hoechst. Capn15 seems to be enriched in Purkinje cells in the cerebellum. Scale bar is 100μm. GL, granular layer; ML, molecular layer.

As homozygous disruption of *Capn15* leads to abnormal eye development (Zha, et al., 2020), we examined Capn15 staining in the eye, although in this case we focused on an earlier stage of development, P3. At this stage, X-gal staining is mainly restricted to the retinal ganglion cell layer (Zha, et al., 2020). To confirm expression of Capn15 in retinal ganglion cells, we examined co-localization of β-gal with a marker for retinal ganglion cells, Brn3a (Nadal-Nicolas et al., 2009). As shown in Fig.9, the two markers co-localized (79 ± 5 of β-gal cells colocalized with Brn3a) but only about half of the retinal ganglion cells contained β-gal (44 ± 2 Brn3a neurons co-localized with β-gal). We also examined co-localization of β-gal with ChAT as a fraction of retinal ganglion cells that do not contain Brn3a are displaced cholinergic amacrine cells (Jeon et al., 1998). Similar to Brn3a staining, about half of the ChAT labelled neurons co-localized with β-gal (62 ± 5). Thus, Capn15 is expressed mainly in the retinal ganglion cell layer in a subset of Brn3a+ and ChAT+ cells.

**Figure 9.**
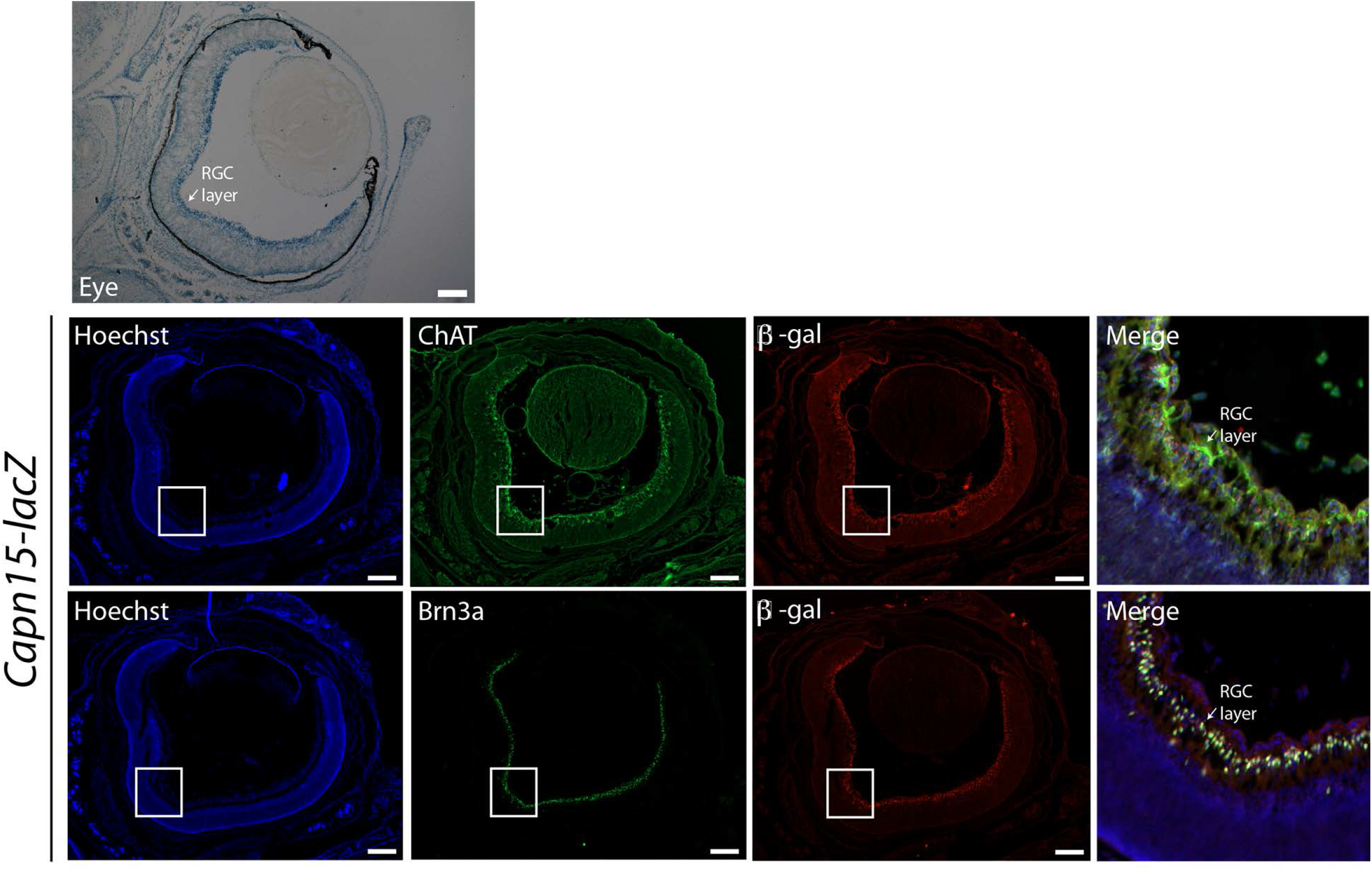
*Capn15* distribution in the eye. *Top*, X-gal staining of coronal sections from P3 *Capn15^(lacZ-Neo)^* heterozygous mice eyes are shown. Scale bar is 200 μm. RGC, retinal ganglion cells. *Bottom*, The distribution of a retinal ganglion cell marker (Brn3a) or that of cholinergic amacrine cell marker (ChAT) is shown along with that of β-galacatosidase (β-gal) in sections from adult *Capn15-lacZ* brains. The chicken polyclonal antibody directed against β-gal and either the goat polyclonal antibody directed against Brn3a or the goat polyclonal antibody directed against ChAT were revealed by using a goat anti-chicken secondary antibody conjugated to Alexa Fluor 594 and a donkey anti-goat IgG secondary antibody conjugated to Alexa Fluor 488 respectively. Capn15 is expressed in the retinal ganglion cell layer in a subset of Brn3a+ and ChAT+ cells. Scale bar is 200 μm.

## Discussion

We have previously described *Capn15* KO mice and found that they had a lower body weight and striking ocular anomalies when compared to their WT littermates (Zha, et al., 2020). In this paper, we further characterize these mice using MR imaging and localization of Capn15 expression in the adult. The loss of Capn15 protein leads to smaller brains and specific changes in the thalamus, amygdala and hippocampal subfields. Capn15 is expressed in a variety of brain regions in the adult including excitatory neurons linked to plasticity changes involved in learning and memory.

The average weight of a whole *Capn15* KO mouse was 11% smaller compared to its WT littermate (Zha, et al., 2020). MRI results showed that the *Capn15* KO brain was 14% smaller in volume compared to the WT animals and the weight of the brain was similarly reduced. One human individual with a biallelic variant CAPN15 had clinical microencephaly (Zha, et al., 2020) as did a patient with a likely loss of function of CAPN15 (Mor-Shaked et al., 2021), consistent with the mouse recapitulating a loss of brain volume also seen in humans. The total change in brain volume was similar to the 11% weight loss, although there is no correlation between body and brain weight in mice (Spring et al., 2007). The brain makes up only a small fraction of the weight of the mouse, so the decrease in brain volume cannot explain the overall weight difference. The loss of whole animal weight and brain volume are likely to be two independent aspects of the *Capn15* KO phenotype.

Certain brain areas such as subfields of the hippocampus and thalamus were still significantly smaller in the *Capn15* KO mice, even when taking the 14% global brain volume change into account. On the other hand, the amygdala was significantly larger in *Capn15* KO mice after scaling. This indicates that either this region is relatively spared or that amygdala volume increases in the KO mouse. It should be noted that only viable *Capn15* KOs were examined and, based on Mendelian ratios, only 50% of *Capn15* KO animals survive to weaning. Thus, it is possible that the mice that did not survive to weaning had more severe decreases in brain structure volume.

We were not able to associate decreases in brain volume with a decreased neuronal number (using Nissl staining) in Purkinje neurons or CA1 neurons, although we did not have the power to detect changes in cell number in the same range as the decrease in brain volume. Brain volume derived from MRI measurement can also be due to many factors other than cell number. In particular, neuronal cell size, astrocyte size and number, as well as synaptic and axonal density and axonal volume fraction can mediate volume changes (Lerch J. P. et al., 2011). It will be interesting in the future to determine more precisely the cause of the overall volume change in these mice.

There does not appear to be a relationship between where Capn15 is expressed in the mature brain and the regions mostly affected by Capn15 loss during development. Thus, our results are consistent with an important role for Capn15 early in brain development that leads to lasting deficits in brain volume in the adult, as opposed to an ongoing requirement for Capn15 for survival. The continued expression of Capn15 in brain areas involved in plasticity is consistent with a possible later role in plasticity, as has been seen in *Aplysia* (Hu, et al., 2017a). This may be similar to Capn2 that plays an important role in early development and a later role in plasticity in the adult (Dutt, et al., 2006;Liu, et al., 2016).

A major unresolved issue is the localization of Capn15 in the dentate gyrus molecular layer. This diffuse staining was not seen in other brain regions, such as the cortex or amygdala. The immunostaining showed much smaller puncta with the antibody to β-gal and the X-gal staining took much longer to saturate, suggesting lower expression in this region. However, the staining did not seem to colocalize with astrocytes, microglia, inhibitory neurons or Hoechst nuclear staining. We do not know the source of this staining.

### Conclusion

Removal of *Capn15* in mice results in smaller brains with larger changes in the thalamus and sub-regions of the hippocampus, although the reasons underlying the change in the size of the brain are still unresolved. In the adult mice, Capn15 expression remains in a subset of excitatory neurons, particularly neurons implicated in plastic changes. The *Capn15* cKO mouse should be a good model for exploring the role of Capn15 in plasticity since its brain is of normal size and lacks the eye anomalies present in the complete KO mouse.

## Acknowledgments

This work was supported by CIHR grant MOP 340328 to WSS. The authors would like to thank Mireille Bouchard-Levasseur for excellent technical assistance in quantification of the transgenic mice eye phenotype and Dr. Len Levine for advice concerning the eye phenotype

